# DoggifAI: a transformer based approach for antibody caninisation

**DOI:** 10.1101/2025.05.28.656573

**Authors:** Dominik Grabarczyk, Mikołaj Kocikowski, Maciej Parys, Douglas R. Houston, Ted Hupp, Javier Antonio Alfaro, Shay B. Cohen

**Author notes:** Corresponding author. E-mail address (D. Grabarczyk). These authors contributed equally to this work (Dominik Grabarczyk, Mikołaj Kocikowski). These authors co-directed this work equally (Javier Antonio Alfaro, Shay B. Cohen).

## Abstract

Antibody translation across species offers a compelling strategy to extend the vast and expensive investments in human therapeutic antibodies to veterinary oncology, with applications in both veterinary medicine and comparative oncology.

While precise, low-immunogenic treatments are essential for canine cancer care, traditional species conversion methods rely on ad hoc bioinformatics modifications. These methods often implicitly decouple the framework (FR) and complementarity-determining regions (CDRs), ignoring how structural changes in FRs can affect the conformation and function of CDRs. This can compromise binding specificity and require costly high-throughput *in vitro* screening.

To address this, we present DoggifAI, a transformer model that translates non-canine antibody sequences into canine ones by generating species-appropriate framework regions (FRs) based on desired CDRs. This allows the model to better preserve structural compatibility between FRs and CDRs. The model is pretrained in a T5-style text-to-text denoising task on a large multispecies antibody dataset, which allows further finetuning on a much smaller species-specific dataset.

DoggifAI generates highly canine-like antibodies and shows promising results in preserving binding specificity. To support further progress in this field, we also release a curated dataset of over 430,000 unique canine antibody chain sequences, significantly expanding the public sequence repertoire.

Graphical Abstract

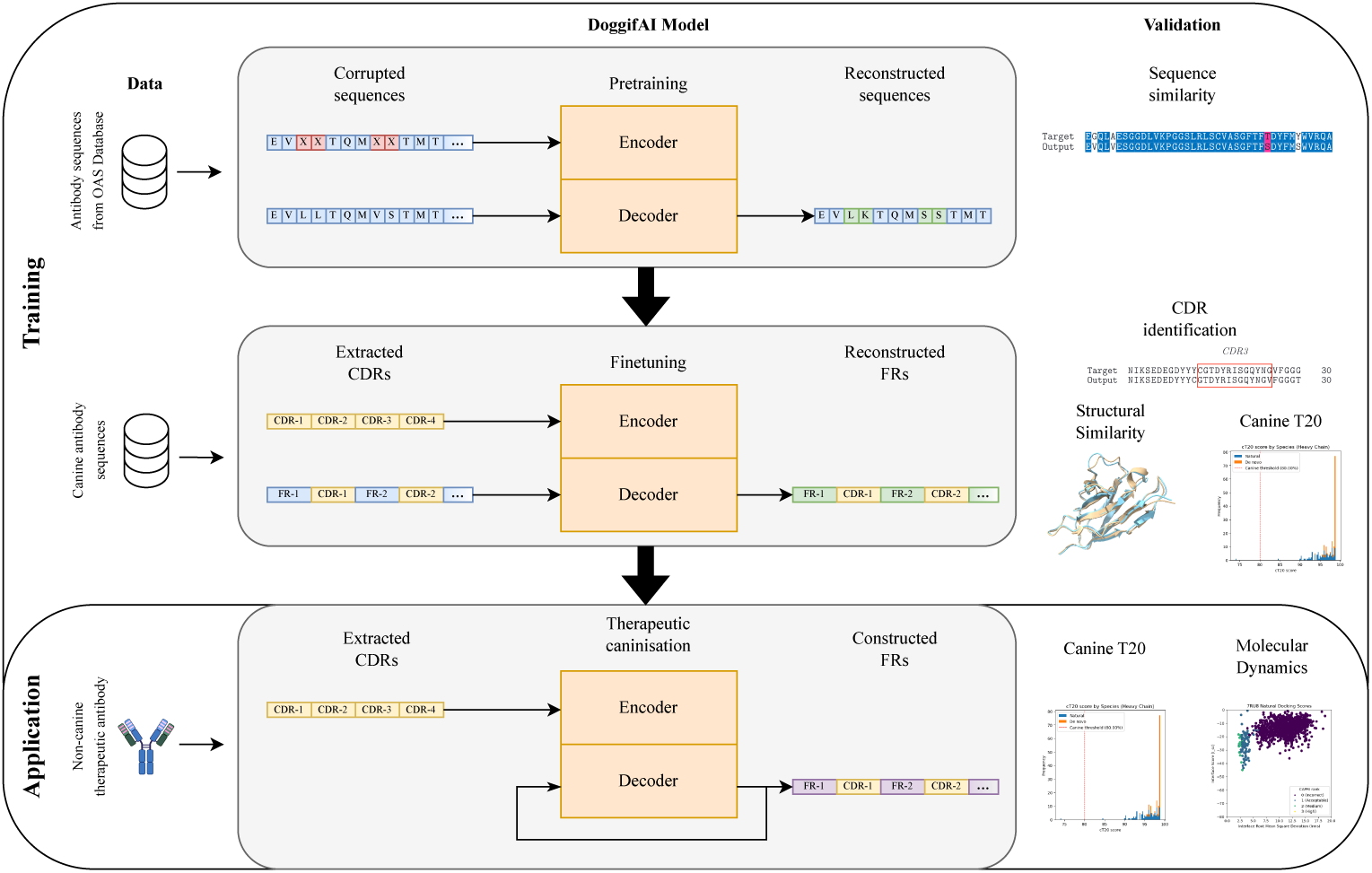

**Highlights:**

- We show that transformer models are capable of generating plausible antibody framework regions based on CDRs
- We show that resulting framework regions are highly recognisable as coming from the desired species
- We show promising results for the retention of binding specificity when translating antibody sequences in this way
- We release a large, high-quality dataset of canine antibody sequences to support future research

## 1. Introduction

Substantial investments in human antibody therapeutics have yielded molecules with finely tuned binding characteristics and minimized immunogenicity, yet their direct application to veterinary oncology is challenged by species-specific variations in antibody frameworks and differences in immunology. Given the significant resources dedicated to human antibody (Ab) development, adapting these molecules for canine applications is both a logical and potentially cost-effective strategy. Conventional approaches to ‘caninisation’ of human or murine antibodies often involve ad-hoc modifications that insufficiently consider the structural nuances of complementarity-determining regions (CDRs), which are essential for antigen binding.

Antibody-based therapies have achieved notable clinical success, demonstrating efficacy in managing viral infections [11, 33], diverse malignancies [36], and autoimmune disorders [10]. An antibody of the IgG class - one most relevant to the therapeutic application - consists of two identical chains. Each chain can be divided into two regions. A ‘fragment crystallisable’ (Fc) region is characterized by a relatively conserved amino acid sequence and responsible for the interaction with other elements of the immune system, such as phagocyte cells and complement proteins. The ‘fragment antigen-binding’ (Fab) region is diverse, and responsible for specific binding to a single target antigen. Fab contains one ‘constant’ and one ‘variable’ domain, with the latter consisting of four framework regions (FR) and three CDRs. The CDRs directly define the specificity of the Ab to an antigen, which is enabled by the diversity of billions possible sequence configurations, and resultant structural diversity [4]. These CDRs are surrounded by FRs which provide a structural scaffold, crucial for the proper folding of the Ab into a functional protein. A schematic overview of these regions can be seen in Figure 1.

**Figure 1:**
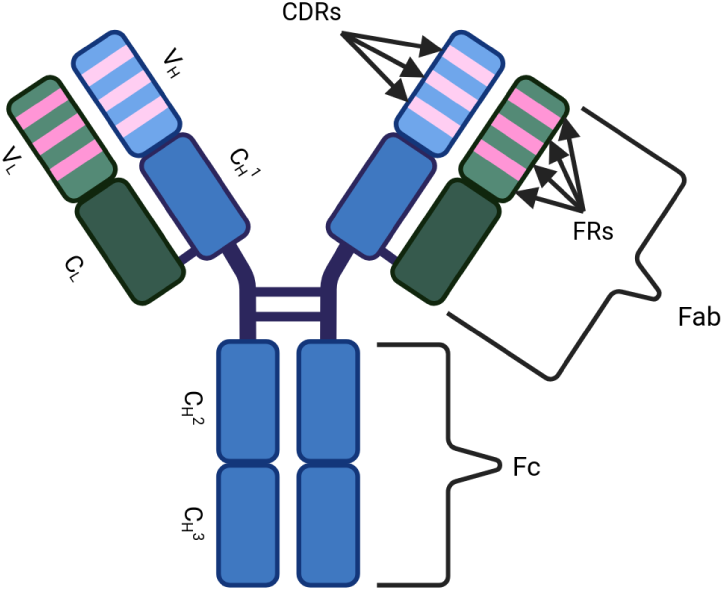
Schematic overview of the structure of an antibody. Fab stands for fragment antibody binding, Fc for fragment crystallisable, *V_H_* for variable domain of the heavy chain, *C_H_*for constant domain of the heavy chain, *V_L_* for variable domain of the light chain, *C_H_* for constant domain of the light chain, CDR for complimentarity determining region and FR for framework region. The pink and purple lines correspond to the CDRs of the heavy and light chain respectively.

The high diversity of antibodies allows massive datasets of antibodies to be collected. These in turn make antibody design a very suitable problem to address with artificial intelligence (AI), which tends to perform well in problems with abundant training data. Many AI tools have been developed for antibody design [19]. For example, some tools seek to replace or augment current experimental antibody development processes by generating CDRs, often focusing specifically on the highly variable CDR-H3 region [34, 25]. Aside from generative tools, many tools have been developed which evaluate Ab viability as therapeutics, for example by predicting their solubility or immunogenicity [31, 21].

Antibodies exhibit enough sequence conservation to be non-immunogenic within a species, even in their high-diversity regions. However, conserved motifs vary by species, meaning that antibodies from a species different from that of the host would be identified as foreign. For example, a mouse antibody injected into a human would bind to the same targets, but cause an immune reaction at the same time. Such reaction poses danger to the patient and reduces the treatment effectiveness.

The ability to translate antibodies between species such as dog and human offers several benefits. First, many cancers in dogs closely resemble those in humans and often involve conserved protein antigens [20]. Translating antibodies between the two species supports comparative oncology and opens new angles for studying cancer immunotherapy-potentially helping to solve the riddle of human cancer through insights gained in canine models. Second, many antibodies are developed in rodents, especially using the mouse hybridoma technology, for *in vitro* research on dog targets. A method to ‘caninise’ these antibodies would allow their clinical application, making the development process more practical and worthwhile, especially given the limited funding in this area. Finally, while a single pathogen can infect multiple species, most clinical trials focus on human therapeutics. Being able to adapt existing antibody treatments to animals would support veterinary medicine in the fight against infectious diseases, especially for dogs.

Approaches to translating protein languages between species, particularly for Abs, are lacking. For example, in one popular translation method, called CDR grafting, the desired CDRs are surrounded by FRs from the target species which exhibit the highest homology to the original Ab’s FRs [24]. However, this method does not take into account the CDR sequences for which the FRs are selected, often leading to deformations in the CDRs and consequently decreased Ab specificity [38]. A related but more involved method called Framework Shuffling involves combining different FRs from an antibody library of the target species with the desired CDRs and using high-throughput screening methods to find plausible candidates [12]. This somewhat alleviates the issues of not tailoring the FRs to the specific CDRs, but requires expensive screening. Both these methods also rely on having big and diverse libraries of Abs for the target species, which may not be the case for less studied species.

We propose an AI model, DoggifAI, for Ab translation which leverages the context provided by CDRs to generate better FR sequences. To address the challenges posed by species for which large Ab databases do not exist, we use T5-style semi-supervised pretraining [30] on a large multi-species Ab database, where the model needs to reconstruct small corruptions in an Ab in a text-to-text manner. This allows the model to learn general structures present in Abs. The model is then fine-tuned on a smaller dataset of canine Abs, where it is tasked to reconstruct the FRs based on given CDRs. By using a transformer architecture, the model can use the context provided by the CDRs to inform its choices for the FRs. We demonstrate our approach is able to generate diverse but highly canine sequences which score well on the reconstruction error. Furthermore, we show promising results for the binding affinity of the resulting Abs when tested *in silico*.

## 2. Materials and Methods

The following section outlines the architecture of the DoggifAI model (subsection 2.1), the data used to train it (subsection 2.2), the training methodology (subsection 2.3) and the validation methods (subsection 2.4)

### 2.1. Transformer model

The model uses a transformer architecture as proposed by *Vaswani et al.*[37]. The architecture sketch can be seen in Figure 2. The model is trained using a T5 style training regimen [30], specifically by pretraining on a sequence-to-sequence denoising task followed by fine-tuning on the sequence construction task. An example of the pretraining and finetuning data can be found in Figure 3. Four different model sizes are tested, their dimensions can be found in Table 1. The hyperparameters used can be found in Appendix B.

**Figure 2:**
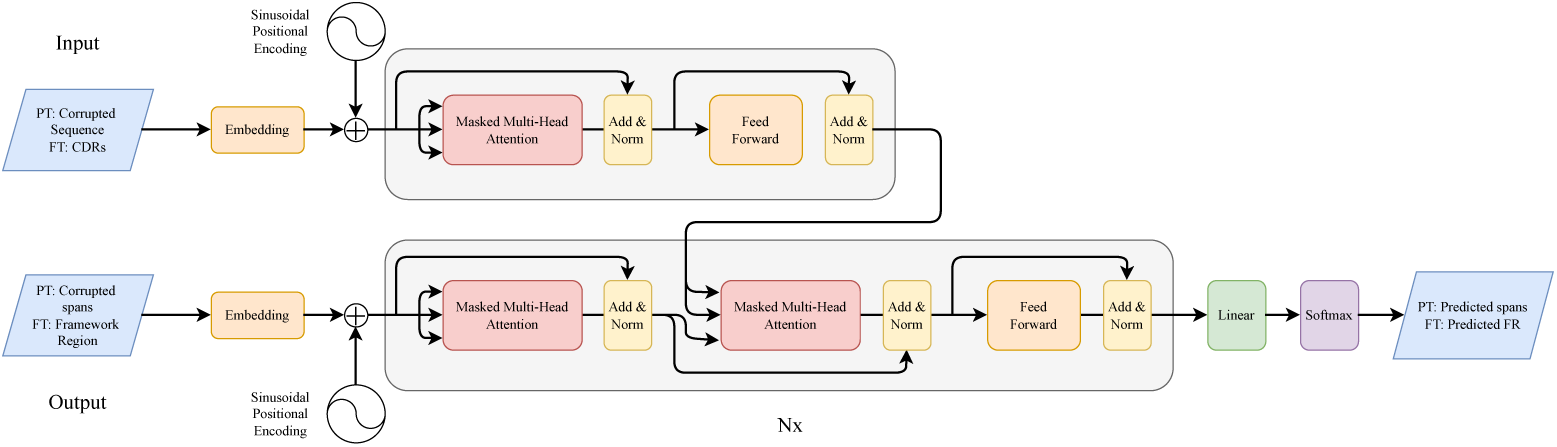
DoggifAI Transformer Model Architecture. PT denotes variables during pretraining and FT denotes variables during finetuning

**Figure 3:**
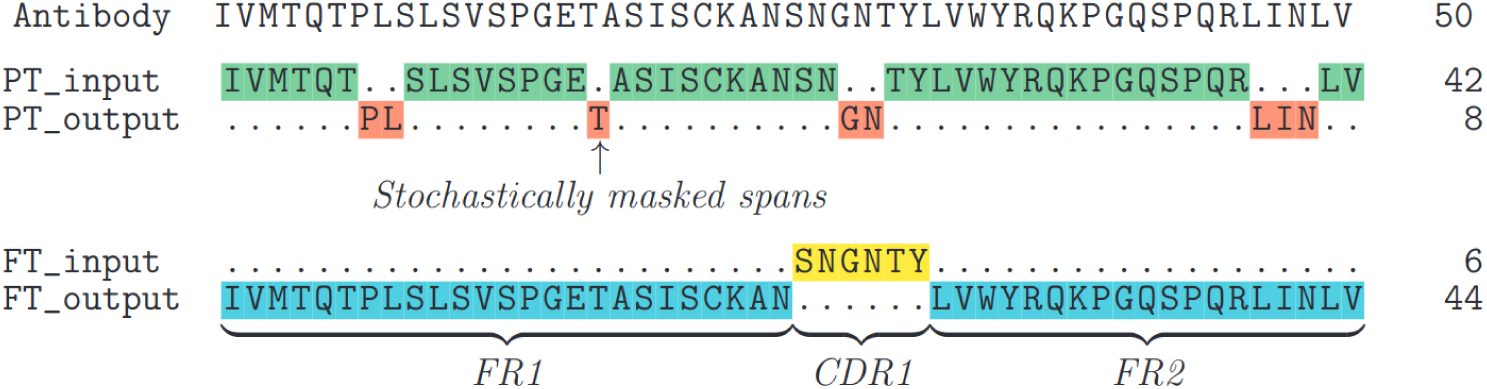
First 50 residues of an example antibody and how it is processed into the pretraining (PT) and finetuning (FT) data. In the pretraining setting the input is created by randomly masking short spans throughout the sequences. These masked spans are used as the target output. In the finetuning setting the CDRs are extracted to serve as model input, and the framework regions are extracted to serve as target output.

**Table 1:** Overview of the tested model sizes. *N* stands for the number of layers, *h* for the number of attention heads, *d_model_* for the dimensions of the hidden layers and *d_ff_*for the dimension of the feed forward layer

| Name | $N$ | $h$ | $d_{model}$ | $d_{ff}$ |
| --- | --- | --- | --- | --- |
| Tiny | 2 | 4 | 128 | 2048 |
| Small | 4 | 4 | 128 | 2048 |
| Medium | 4 | 4 | 256 | 2048 |
| Large | 8 | 8 | 512 | 2048 |

Two strategies are implemented for sampling the output sequences: greedy sampling and beam search. Greedy sampling is implemented by using a top k sampler with *k* = 1. This method is used during training and validation due to its low computational cost compared to alternatives. In this sampling method the model generates the sequence from left to right, at every step picking the element with the highest likelihood. Beam search is often used to generate more accurate samples at test and inference time. In this set up, rather than adding the highest likelihood token to the sequence, the model keeps track of the top *n* highest likelihood sequences, extending them greedily. This allows the model to better avoid local likelihood maxima, where a high likelihood token does not have any reasonable follow up tokens.

### 2.2. Datasets

Two separate datasets are used in this project for training. Firstly, a large but non-specific dataset of unpaired antibody sequences from different sequencing projects and species used for pretraining coming from the Observed Antibody Space database (see subsubsection 2.2.1). Secondly, a smaller canine dataset of unpaired antibody sequences used for finetuning (see subsubsection 2.2.2)

#### 2.2.1. Observed Antibody Space Dataset

The Observed Antibody Space (OAS) dataset is constructed from the unpaired heavy and light chain sequences data available on the OAS portal [28]. These sequences come from a wide variety of sequencing projects and are diverse in terms of the species, diseases and isotypes. The majority of the sequences are human (92%) and murine (7%) with the remainder split between rabbit, rat and rhesus monkey sequences. 85% of the sequences are heavy chains and 15% light chains. The average length of the variable domains is 111 amino acids. Duplicate entries were then removed, as well as sequences which are smaller or larger than sequences found in the canine dataset.

The resulting dataset comprises over 2 billion sequences, encompassing mixed heavy, light kappa, and light lambda chains. As this volume exceeds the computational infrastructure currently available at our academic institute, a random subset of 10 million sequences was selected to form the final dataset, hereafter referred to as the OAS dataset.

#### 2.2.2. Canine antibody Dataset

The canine dataset is based on a phage scFv library by *Lisowska et al.* [23] which was generated from spleen tissue samples from multiple dogs. Briefly, mRNA was previously extracted, pooled, and amplified using RACE-PCR. Degenerate primers were used to clone heavy– and light-chain sequences into plasmids. The light and heavy chain holding plasmid libraries obtained in this way were then combined to construct a phage display library of single-chain variable (scFv) antibody fragments. Three separate libraries-heavy chain, light chain, and scFv-were subjected to long-read, high-fidelity sequencing on a PacBio platform.

The resulting sequences were pooled and analysed using in-house software and manual curation, producing a database of unpaired heavy– and light-chain sequences. Antibody encoding sequences were identified based on sequence tags, translated into amino acid sequences, and de-replicated. Sequences missing any structural frameworks (FR) or complementarity determining regions (CDR), or containing ambiguous amino acids, were excluded, except for those with truncated FR1 or FR4 regions (since these are the beginning and end of the chain). Additionally, sequences classified as ‘unproductive’ by IMGT High-V-Quest [9] were removed.

Unproductive sequences were defined as those that were unlikely to yield functional antibodies due to disrupted reading frames, premature stop codons, or unsuccessful V(D)J recombination. Although this filter may reduce repertoire diversity, especially given reliance on known germline sequences, such filtering was necessary to ensure a high-confidence dataset for training an AI model for functional antibody generation.

The final dataset consisted of 272,535 unique heavy chain sequences, 79,545 light kappa chains, and 81,881 light lambda chains. The dataset exhibited high CDR diversity, particularly in CDR-H3, which is the most rearranged and functionally critical region. After preprocessing, the dataset was refined to a total of 425,284 sequences.

The Ab variable regions were separated into CDRs and FRs using the IGMT numbering scheme and Abnum web-tool [1].

### 2.3. Training

The model is trained in three different setups which are compared and contrasted.

1. **Finetuning on canine dataset only** In this set up, the model is trained only on the final task of constructing the framework regions around the CDRs on the canine dataset.
2. **Pretraining and finetuning on canine dataset** In this set up, the model is trained using a T5 pretraining regimen [30]. The model is first pretrained on the canine dataset in a semi-supervised sequence-to-sequence fashion where random spans in the sequence are corrupted and the model is tasked with restoring the corrupted tokens. Then the model is finetuned, using the same canine dataset, on the final framework region construction task. This method should investigate if it is possible to train twice on the same data in data-sparse environments common in therapeutics design to improve performance and convergence rates.
3. **Pretraining on OAS dataset, finetuning on canine dataset** Here the model is pretained using the same sequence-to-sequence method explained in the previous paragraph on the OAS dataset. This step simulates a more common scenario where pretraining data is larger and more diverse than the small finetuning dataset. It should let the model learn the general structure of antibodies and make the finetuning converge faster and more effectively. Finetuning is again performed on the canine dataset.

During pretraining on average 15% of tokens are corrupted per sequence in spans with an average length of 3 tokens.

### 2.4. Validation

This section outlines the various experiments we performed to validate the DoggifAI *de novo* generated sequences. The validation tasks are performed for a varying number of sequences for each task, as some properties are computationally cheap to validate like comparing the reconstructed FR sequences to the original FR sequences, while others are highly complex like the antibodies’ binding affinity.

#### 2.4.1. Sequence alignment Loss

The generated sequences are compared to the ground truth sequences using global pairwise sequence alignment [39]. We derive two different alignment loss metrics, Hamming Distance Loss *L_h_* and Biological Similarity Loss *L_b_*.

*L_h_* is the minimal number of substitutions and gap insertions needed to match two sequences. While *L_h_* is easily interpretable, in practice, different amino acid substitutions are not equally impactful on the folding and functionality of a protein, as some amino acid pairs are more similar in terms of chemical properties than others. Similarly, a single gap of length *n* in the sequence is generally less impactful than *n* single amino acid gaps.

To address this, we also define a biological similarity loss *L_b_*, where amino acid substitutions are weighted using the PAM30 substitution matrix [13], gap insertions are penalised by *−*11 and extensions to an existing gap are penalised by *−*1.

#### 2.4.2. Canine antibody similarity

In antibody humanisation, the T20 humanness score is often used to quantify the degree to which a *de novo* antibody is human-like [16]. Calculating the T20 score involves performing a protein multiple sequence alignment (MSA) using the Basic Local Alignment Search Tool (BLAST)[6] against a curated database of human antibodies. The alignments are scored based on the percentage of identity matches, i.e. the percentage of amino acids in the sequences which match exactly. The best 20 identity match percentages are then averaged to obtain the T20 score.

We reimplement the T20 scoring method for scoring canine-likeness by constructing framework-only T20 cut-off canine databases from the training and validation data using the methodology from *Gao et al.* [16]. These databases are used to obtain canine T20 scores (cT20).

Another metric we used to assess the quality of the *de novo* Abs is whether the same CDRs are extracted by the CDR extraction algorithm as in the original sequences. Firstly, if the CDR extraction algorithm cannot recognise clear CDRs, that might indicate severe structural flaws in the *de novo* sequences. Furthermore, if CDRs are detected, but differ from the CDRs used as algorithm input it might mean they have been inserted at the wrong location by the model. Therefore, for each of the selected models the number of sequences where CDR extraction failed is reported. CDR extraction failures occur in our model outputs in repetitive, wrongly generated sequences. For the successfully extracted CDRs the average *L_h_* between the extracted *de novo* CDRs and the orignal CDRs is reported.

#### 2.4.3. Structural analysis

Alphafold3 [3] is used to make 3D folding predictions of the sequences. The 3D structures of the original sequences and the *de novo* sequences can then be compared to assess structural similarity using Template Modelling (TM) scores [40] and Root Mean Square Distance (RMSD) between the structures. TM scoring differs from RMSD in two ways. Firstly, it is normalised to a (0, 1] scale, which makes it possible to compare scores of proteins with different lengths. Secondly, it measures alignment between substructures making it less sensitive to local deformations.

#### 2.4.4. Caninisation of Existing Antibodies

DoggifAI is designed to create novel canine FRs that are adapted to CDRs from other species while preserving antibody specificity. Specifically, FRs which preserve the specificity of the antibody.

We next focus on the adaptation of five therapeutic antibodies that have been experimentally modelled in complex with their ligands – in regards to newly designed sequences binding the same target. Four of these are proteins targeted for cancer immunotherapy, specifically Programmed cell death protein 1 (PD-1) [42], Programmed death-ligand 1 (PDL-1) [22], Epidermal growth factor receptor (EGFR) [17] and Cytotoxic T-lymphocyte associated protein 4 (CTLA-4) [15]. The last is an antibody therapeutic targeting the SARS-CoV2 Spike Protein (SARS-CoV2 S) [41]. All of these have been retreived from the Protein Data Bank (PDB) [8].

*De novo* sequences were generated using DoggifAI for both chains of each of the five therapeutics. These paired *de novo* sequences were folded as complex using Alphafold3. These folded structures were then relaxed using a molecular dynamics simulation using GROMACS [2]. Finally, they were docked to the original ligand using Rosetta Snugdock protocol [35] and scored based on its Interface Score (I sc) [5], Interface Root Mean Square Deviation (Irms) and CAPRI rank. To obtain a baseline for high docking score each original antibody is also docked to its ligand. To obtain a baseline low docking score, each original antibody is docked with the ligands of each of the remaining antibodies.

## 3. Results

### 3.1. Model Selection

The models are pretrained for 100 epochs and the best epoch is chosen based on the lowest biological similarity loss on the validation set. These best epochs as well as a randomly initialised model are finetuned on the FR reconstruction task for a further 100 epochs. The summary of the results for the finetuning tasks can be seen in Table 2.

**Table 2:** Results for the best epoch of the trained models based on their validation set performance. *L_ce_* stands for crossentropy loss, *L_b_* stands for biological similarity loss and *L_h_* stands for the hamming distance. Model sizes can be found in Table 1.

| Model | Epoch | $L_{ce}$ | $L_b$ | $L_h$ |
| --- | --- | --- | --- | --- |
| Tiny no PT | 42 | 0.3719 | 174.96 | 17.56 |
| Tiny FT | 47 | 0.3966 | 181.63 | 17.81 |
| Tiny OAS FT | 19 | 0.2947 | 91.42 | 8.91 |
| Small no PT | 12 | 0.2908 | 114.09 | 11.54 |
| Small FT | 14 | 0.2491 | 95.52 | 9.39 |
| Small OAS FT | 38 | 0.3097 | 83.43 | 8.33 |
| Medium no PT | 1 | 0.3752 | 133.25 | 13.44 |
| Medium FT | 1 | 0.3700 | 266.53 | 27.31 |
| Medium OAS FT | 23 | 0.2504 | 120.63 | 11.74 |
| Large no PT | 12 | **0.2366** | 107.31 | 10.76 |
| Large FT | 28 | 0.2453 | 108.54 | 10.97 |
| Large OAS FT | 14 | 0.2460 | **74.90** | **7.65** |

The results demonstrate that pretraining on a large and varied dataset yields significant improvements over the other model settings. The utility of reusing finetuning data for pretraining seems limited, and can even increase the loss compared to not pretraining at all.

It should be noted that the sequence alignment losses plateau within approximately five epochs for all models, regardless of the trajectory of the cross-entropy loss. That is to say, the sequence alignment losses remain fairly static, even while cross-entropy loss is still decreasing and after the cross-entropy loss starts increasing due to overfitting.

In the next part the four OAS pretrained models are compared on the test set.

### 3.2. Sequence analysis

The four OAS pretrained models generate 1000 sequences based on CDRs from the held-out test set. The sequence alignment scores for the greedy and beam search sampling methods are provided in Table 3. While the differences, other than the medium sized model, are modest, the large model performs best. This is in line with the findings of Kaplan et al. [18] for large language models (LLMs). *Kaplan et al.* suggest for a dataset of about 50 million tokens like our canine dataset, a model with about 25 million parameters will perform best, which corresponds to our Large model setting.

**Table 3:** Average biological loss *L_b_* and hamming distance *L_h_* between generated sequences and test sequences for the various pretrained model sizes.

|  | Greedy |  | Beam Search |  |
| --- | --- | --- | --- | --- |
| | $L_h$ | $L_b$ | $L_h$ | $L_b$ |
| Tiny OAS | 9.349 | 98.140 | 10.252 | 107.023 |
| Small OAS | 8.737 | 87.775 | 9.547 | 97.131 |
| Medium OAS | 12.101 | 126.497 | 12.499 | 129.675 |
| Large OAS | **8.126** | **80.693** | 8.751 | 87.743 |

The more computationally expensive beam search leads to performance loss compared to the greedy sampling strategy. Since beam-search is only guaranteed to find sequences with higher probability (i.e. lower cross-entropy loss), this is likely related to the previous finding that minimising crossentropy loss does not always yield better alignments.

The Large OAS greedy outputs are validated further as the best model. We analyse the biological alignment loss further by plotting its per-chain score distribution in Figure 4. We see that the model is performs worst when generating light lambda chains with an average *L_b_* of 115.24, compared to the respective 74.82 and 67.10 of the heavy and light kappa chains. This is not surprising as light lambda chains are relatively under-represented at 19% of total data.

**Figure 4:**
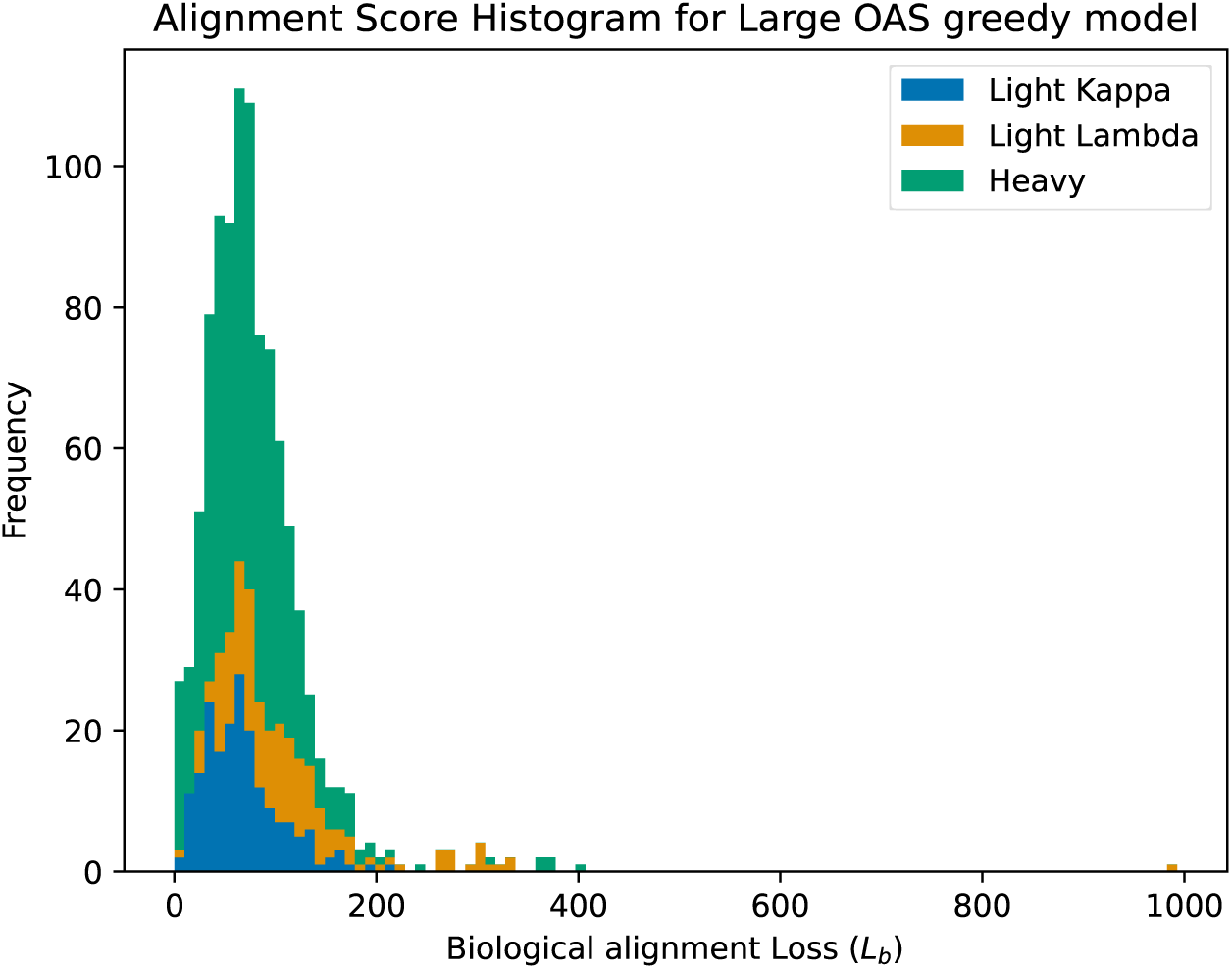
Histogram of the biological losses of the Large OAS greedy model on the test set. The colours represent the 3 chain types.

We also see one very significant outlier. Closer inspection reveals the prediction completely failed and the sequence is gibberish (see Appendix A for the full sequence). It should be noted that such failures do not occur with beam search. However, for the purposes of antibody development trivially unsuitable sequences are not necessarily a problem, as sequences would be tested extensively *in silico* first.

### 3.3. CDR preservation

The 1000 *de novo* sequences from the previous section are split into FRs and CDRs in the same way the original sequences were. In eight cases the extraction failed. The resulting CDRs are compared to the original ones, in nine cases the CDRs did not match. It should be noted that all of those mistakes, both failed extractions and misalignments, are in light lambda chains, which already had been observed to be a weakness of the model. Furthermore, we check if the correct call is made regarding the chain type. In all cases where extraction succeeded the call was made correctly.

### 3.4. Canine T20

For each of the 1000 sequences the cT20 score is calculated. We plot the distribution of the cT20 scores for the *de novo* sequences compared to the distribution of the cT20 scores for the original sequences in Figure 5.

**Figure 5:**
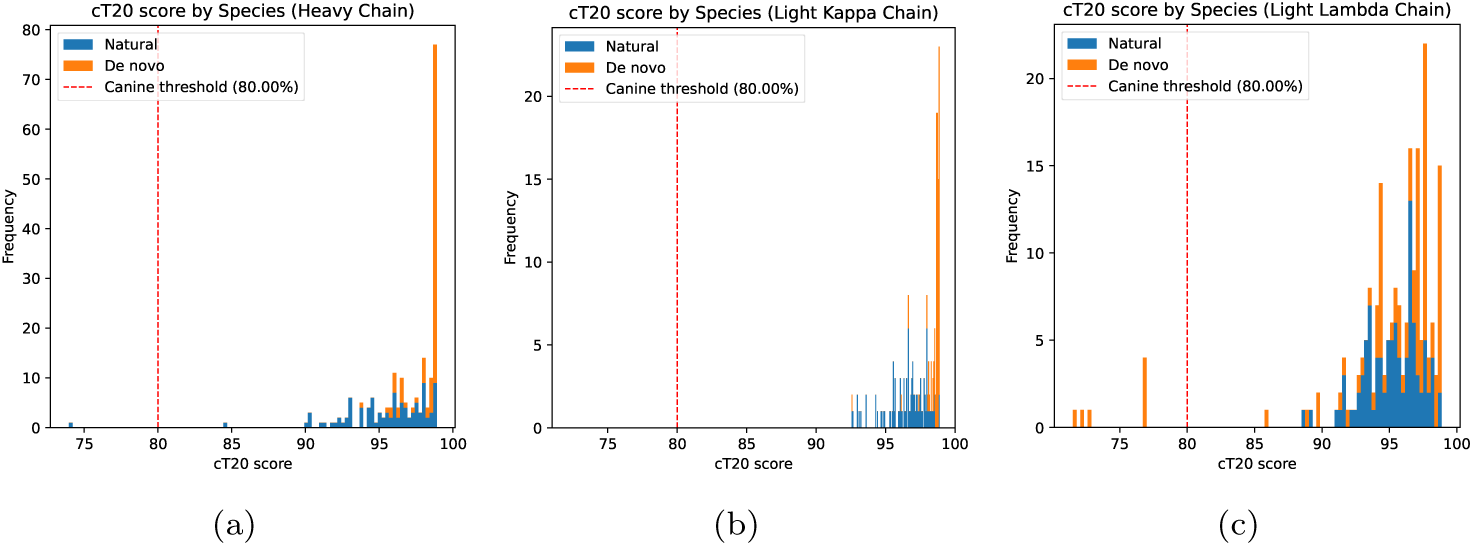
Subfigures (a), (b) and (c) show the cT20 score histograms for the heavy chains, light kappa chains and light lambda chains respectively. The natural sequences are shown in blue and the *de novo* sequences in yellow. The 80% cut-off to consider a sequence canine is shown as red dotted line.

Note that we use a cut-off score of 80 to determine if a sequence is canine. The cT20 scores of the *de novo* sequences, however, are much higher than this in all cases except light lambda chain. In fact most scores are much higher than the natural sequences.

### 3.5. Folding analysis

100 randomly selected *de novo* sequences as well as their natural counterparts are folded using Alphafold3. The average TM-score between the respective natural and *de novo* sequences is 0.91 and the average RMSD is 0.6622 A. A histogram of the TM-scores can be found in Figure 6.

**Figure 6:**
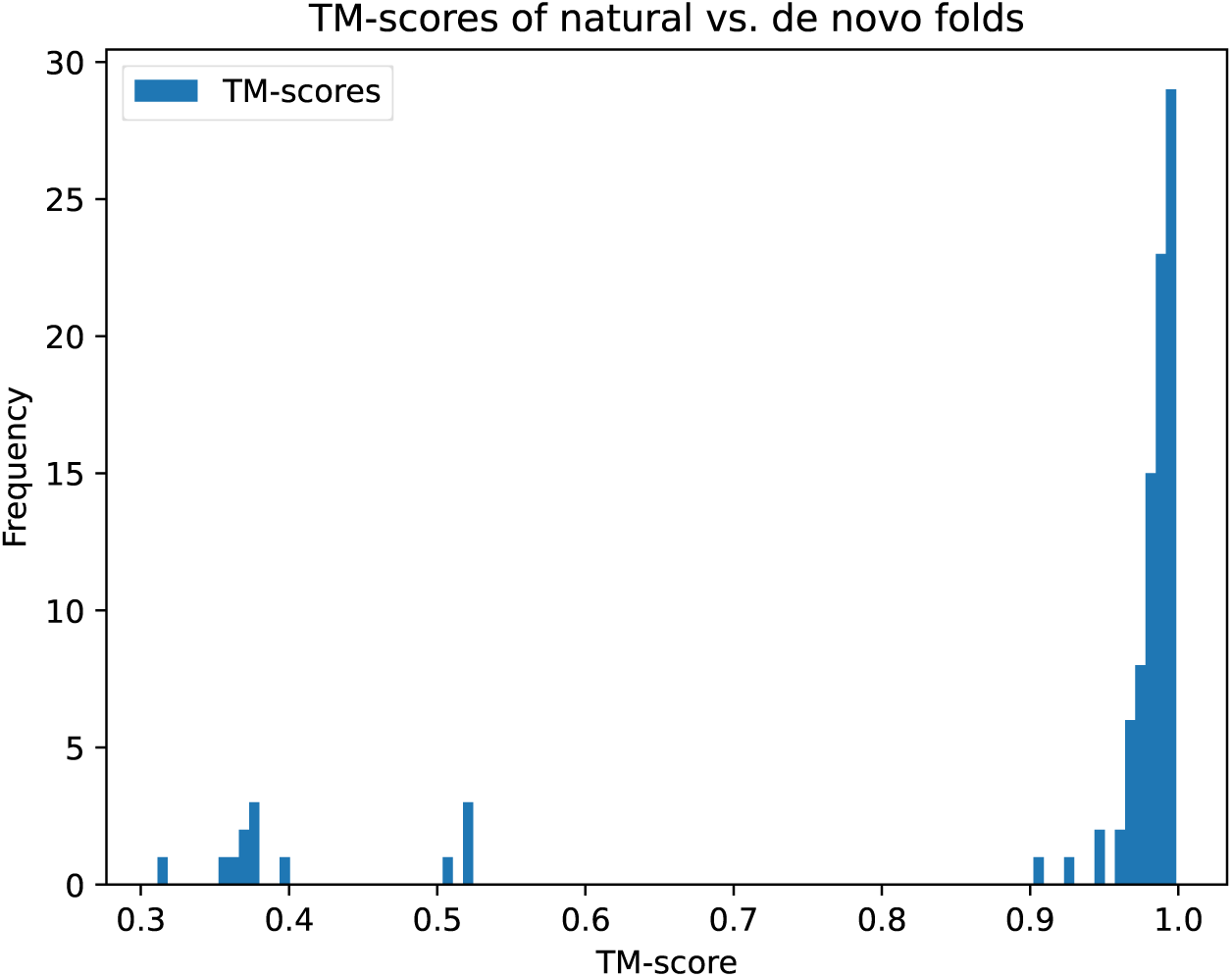
Histogram of the TM-scores between a 100 randomly sampled *de novo* sequences and their naturally occuring counterparts in the test set. A TM-score of 1 indicates a perfect match.

We again observe a sharp peak near the perfect TM-score of 1, and note anomalous failure in generation for a small number of sequences.

### 3.6. Caninisation of Existing Therapeutics

We first extracted the CDRs from both chains of the 5 therapeutic Abs. These were used to generate caninised *de novo* Ab sequences. All caninised *de novo* sequences have a high cT20 score of *>*95, far above the 80 threshold. For each of the 5 ligands we caninised Abs for, we run 6 docking analyses obtaining 1000 decoys per run. Firstly we dock the caninised *de novo* Ab with the intended ligand. Then we dock the original Ab with its intended ligand to obtain a target score to compare *de novo* scores against. Lastly we obtain four negative examples by docking the ligands with the four other natural Abs (e.g. for the PD1 ligand this means docking with the SARS-CoV2 S, CTLA4, EGFR and PDL1 Abs). It should be noted, however, that there is no objective cut-off one can use to decide if a certain score implies a docking, even having our “target” and “negative” scores.

We observe a wide range of results. The best result was found for the SARS-CoV2 S protein. While the score is still worse than the original Ab, it is significantly better than the negative samples. Others, such as EGFR seem to perform similarly to the negative examples. Figure 7 shows the detailed plots of the two aforementioned protein scores, as well as a table of the remaining proteins.

**Figure 7:**
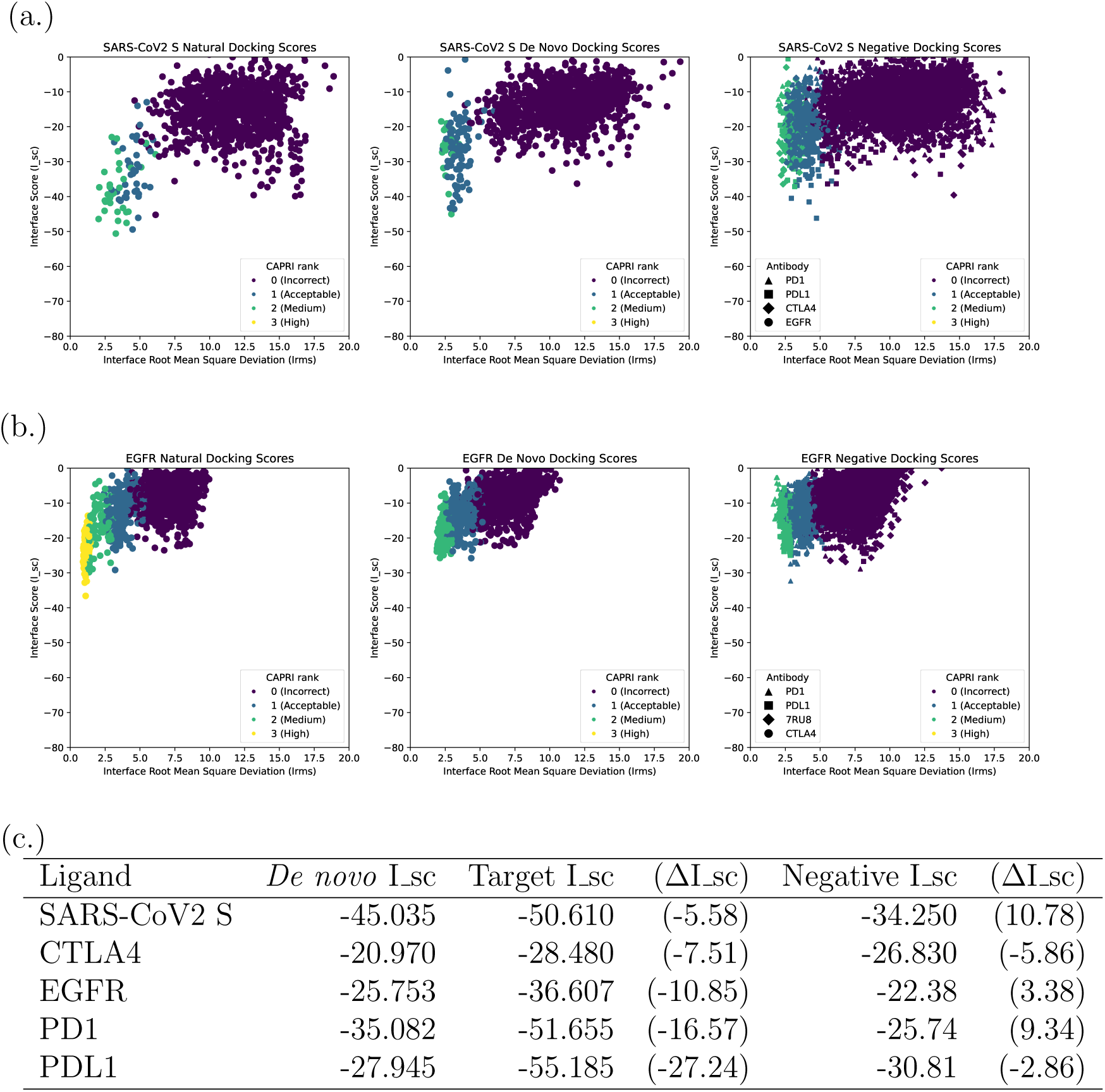
(a.) and (b.) Show the docking results for the SARS-CoV2 spike protein and EGFR respectively. The natural docking score plot shows the results for the ligand with its original Ab. The *de novo* docking score shows the results for the ligand with the *de novo* Ab generated using DoggifAI. The negative docking score plot shows the docking scores for the ligands with the four other Abs targetting different ligands. In all three cases the Interface Score (I sc) is an estimate of the energy of the decoy and the Interface Root Mean Square Deviation (Irms) the distance between the decoy interface region and the original position of the interface region. (c.) A table giving an overview of the docking I sc results. The *de novo* I sc is taken as the lowest I sc found among the highest CAPRI rank decoys of the ligand docked with the *de novo* Ab. The target I sc is taken as the lowest I sc found among the highest CAPRI rank decoys of the ligand docked with its respective Ab. The negative I sc is obtained by first obtaining the lowest I sc among the highest rank decoys for each of the four pairings of the protein ligand with the incorrect natural Ab and then averaging these four scores (e.g. for PD1, the PD1 ligand paired with the SARS-CoV2 Spike glycoprotein Ab, the CTLA4 Ab, the EGFR Ab and PDL1 Ab.). The delta values are obtained by subtracting the respective value from the *de novo* I sc.

As can be seen for example in the results for CTLA4, docking analyses are not a perfect method. For CTLA4, the I sc of the experimentally validated Ab (–28.480) does not significantly differ from the ligand paired with the remaining four Abs (–22.38). This shows that the *in silico* validation is only a preliminary step before more thorough analyses in a lab.

## 4. Discussion

DoggifAI shows that antibody framework regions can be translated to canine sequences using a transformer model guided by CDR context. Unlike heuristic approaches, which typically apply rule-based modifications or rely on sequence similarity without structural awareness, DoggifAI learns patterns from large multispecies data to generate structure-aware outputs. This reduces dependence on high-throughput *in vitro* screening and better preserves binding-relevant features. While validation is currently limited to *in silico* modelling, results suggest strong retention of CDR characteristics, high caninisation, and favourable structural properties. By enabling rapid and cost-effective antibody translation between species, this approach to antibody development can support comparative oncology studies and expand access to antibody-based treatments in veterinary medicine.

In addition, we release a curated dataset of over 430,000 unique canine antibody chain sequences, vastly expanding the publicly available repertoire, which-based on current NCBI Protein database searches-currently includes fewer than 1,000 canine immunoglobulin entries, many of which are incomplete or not true antibody chains.

DoggifAI model is very effective at reconstructing canine Ab sequences with remarkably high cT20 scores, which demonstrates their high canine-likeness.

Slightly lower performance was observed for light lambda chains compared to kappa chains, as shown in Figure 5. This is reflected in a broader score distribution, including some outliers falling below the cT20 threshold of canine likeness. Two factors may explain this discrepancy. First, while light lambda and kappa chains were similarly represented during the canine fine-tuning phase (both lower than heavy chains, which formed approximately 60% of total), lambda chains were under-represented during pre-training on the OAS dataset, comprising only 5% of sequences compared to 10% for kappa. This imbalance resulted from random subsampling of the OAS dataset and could be corrected in future work by rebalancing chain types during training. Second, lambda chains appear to exhibit greater natural sequence variability, as indicated by the broader distribution of native lambda sequences in Figure 5 (blue). As a result, generated lambda chains are expected to show more variation in score. The few sequences generated that fell below the threshold were rare, easily identifiable and would typically be filtered out during *in-silico* validation prior to experimental use.

Our model takes a radically different approach compared to most AI based humanisation techniques, which tend to be centred around learning to score human-likeness first, for example by learning to predict the T20 score of a sequence. Then the resulting models are used to suggest mutations to non-human sequences which would improve the T20 score. For example Sapiens [29] uses attention based deep learning to suggest mutations which would maximise its human-likeness. Hu-mAb [26] uses random forests to obtain a measure for humanness of a sequence, and then uses the model to suggest the fewest changes with the biggest impact on humanness score. A potential problem with this approach is that by optimising only for human-likeness, the suggested mutations might negatively impact other metrics such as therapeutic developability or protein stability. Our model is capable of producing highly canine sequences without being shown explicit canine-likeness metrics. Furthermore, by generating entire FR sequences, our model can explore more of the available design space.

We see promising results in the validation of the therapeutic sequences in particular, where we show our method is capable of creating sequences which retain a high binding affinity to their target while drastically increasing the cT20 score. While we did not obtain high binding affinity prediction for all five of the greedily generated sequences, the advantage of a transformer based method is that we can easily generate hundreds of different candidates to validate by e.g. implementing stochastic sampling approaches. Furthermore, LLMs and transformers benefit greatly from scale, and the necessary components to increase the scale of this project exist. We were able to process only a fraction of the publicly available pretraining data, and further pretraining tasks such as CDR identification can easily be implemented. Alternatively, more broadly pretrained protein language models (PLMs) such as ESM2 [32] could be used instead of our pretraining method. Such PLMs have been shown to outperform immune specific LLMs on immune-system related tasks [14]. While it is costly to use all the data we have available or to use significantly larger models, these costs are insignificant compared to the costs of failed clinical trials which this approach can prevent.

In the context of canine veterinary science, our approach enables rapid and cheap development of experimental antibody therapeutics, provided that antibodies against the target exist-whether originally raised in another species, such as mice, or developed as human antibodies. This is possible where lab-grade antibodies to canine targets were raised for *in vitro* studies or for many important antigens that are targeted in humans and highly conserved across species. For example, certain epitopes of CD20 are shared between dogs and humans, making it a viable target. Notably, CD20 was the first target of an approved antibody therapy for cancer.

Still, for many therapeutically relevant targets, the canine epitope differs from the human one, and no rodent-originating dog-specific Abs exist, leaving no template sequences to caninise. However, *in-silico* generation of human antibodies against novel epitopes is a topic of great interest [7]. Our method could easily be combined with such designed antibodies to generate caninised therapeutics against previously inaccessible epitopes.

The potential of this approach extends beyond veterinary applications and could be applicable to human antibody therapeutics. In the context of comparative oncology, it might be possible to leverage targets with highly conserved sequences and adapt antibodies against them from model animals to humans.

As we are a computational group, all our validation was performed *in silico*. Predicting binding affinity between Abs and antigens remains an active area of research [27]. Wet-lab experiments are a necessary next step to validate our findings. These experiments can then inform further tuning of the model to adapt it to clinical practice. For instance, testing could reveal inconsistency at generating developable Abs. However, the model’s T5-style pretraining makes it well suited for extension: new tasks can be introduced to address such issues without revisions to the core architecture.

DoggifAI offers a flexible, data-efficient approach to species-specific antibody design, with potential to accelerate veterinary therapeutics and comparative cancer research. To support this further, we release a large, high-quality canine antibody dataset as a community resource.

## 5. CRediT authorship contribution statement

**Dominik Grabarczyk:** Writing – original draft, Visualization, Validation, Software, Methodology, Investigation, Formal analysis, Data curation, Conceptualization **Mikołaj Kocikowski:** Writing – original draft, Validation, Resources, Investigation, Data curation, Conceptualization **Maciej Parys:** Writing – review & editing **Douglas R. Houston:** Writing – review & editing, Validation **Ted Hupp:** Writing – review & editing, Resources, Data curation **Javier Antonio Alfaro:** Writing – review & editing, Supervision, Project administration, Conceptualization **Shay B. Cohen:** Writing – review & editing, Supervision, Resources, Project administration, Conceptualization

## 7. Data availability

The canine dataset used to train the model, including the files for light kappa, light lambda, and heavy immunoglobulin chains as well as the trained Large OAS model, is available from: https://doi.org/10.5281/zenodo. 15125375.

The OAS dataset can be provided upon request.

The code is available on Github (https://github.com/Dominko/DoggifAI)

## 8. Acknowledgements

This work was supported by the United Kingdom Research and Innovation (grant EP/S02431X/1), UKRI Centre for Doctoral Training in Biomedical AI at the University of Edinburgh, School of Informatics and the Royal Academy of Engineering UK Intelligence Community Postdoctoral Research fellowship grant number (ICRF2122-5-133). We appreciate the provision of compute resources through the Baskerville cluster (University of Birmingham / EPSRC).

## Appendix A. Miscreated sequence

Below you can find the miscreated light lambda sequence made by the Large OAS greedy model. Identical amino acids are purple, similar ones are pink.

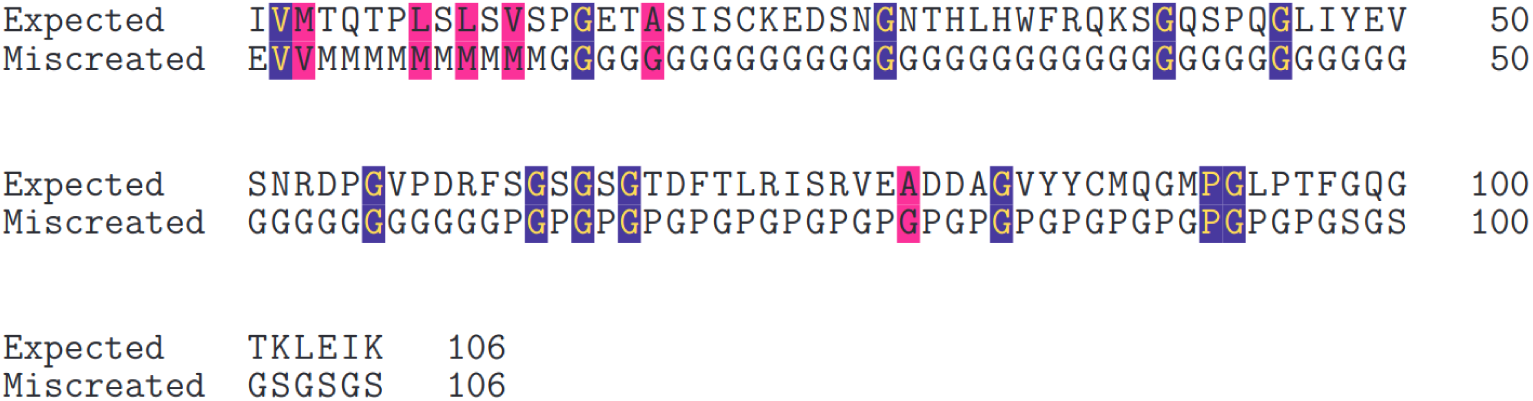

## Appendix B. Hyperparameters

Hyperparameters are kept mostly consistent between the model sizes except for where computational requirements did not allow for it. The models are trained using an ADAM optimiser with a learning rate of 1*e −* 4. A dropout rate of 0.1 is used. The batch size used is 258. No regularisers are used. The models are trained for 100 epochs, except for the large model pretraining on OAS data, which is limited to 10 epochs due to computational resource limitations.

